# Unsupervised source separation and clustering of soundscape recordings predicts avian richness in hyperdiverse systems

**DOI:** 10.64898/2026.10.13.758826

**Authors:** W. Alexander Hopping, Sidd Srinivasan, Oliver C Metcalf, Nárgila Gomes de Moura, Alexander C Lees, Erika Berenguer, Liana Chesini Rossi, Cássio Alencar Nunes, Jos Barlow

## Abstract

1. Passive Acoustic Monitoring (PAM) has emerged as a leading approach for biodiversity research, but analysing the vast quantities of data produced by this method remains challenging. Acoustic indices are often employed to estimate species richness in soundscape recordings, but their generalisability and overall efficacy in speciose regions has increasingly been challenged. An alternative, multi-species classification models, still suffer from performance limitations, particularly in the tropics.
2. We introduce an unsupervised approach that estimates avian species richness using an integrated source separation model trained on birds (BirdMixIT), a leading bird vocalization embedding model (Perch 2.0), and hierarchical agglomerative clustering. We count the clusters of embeddings generated from separated channels to estimate the species richness in recordings at differing timescales.
3. We test the generalisability and predictive performance of our approach, “source-clustering”, using a large combined dataset featuring 103 total hours of manually-annotated soundscape recordings from the Amazon rainforest in Peru and Brazil. To compare the performance of source-clustering to existing approaches, we estimated species richness in the same recordings with both traditional acoustic indices and Perch classifications at various confidence thresholds.
4. Our source-clustering method performed consistently well between our separate datasets. Source-clustering performed better for predicting avian species richness than traditional acoustic indices at all levels, and was complementary with Perch, performing similarly overall to its best settings combinations, though it was more robust for short recordings, and should be more resilient in locations with less training data. Source-clustering generally predicted species richness better than the best-performing Perch settings at the 15-second, 1-minute, and 15-minute temporal resolutions, while most settings of Perch performed better at the 1-hour resolution.
5. Source-clustering can complement or exceed the performance of presently established methods for estimating avian species richness from soundscape recordings, namely traditional acoustic indices and the out-of-box versions of leading automated identification modes. This is particularly true at narrow temporal resolutions, in systems where training data is scarce, and where species density is high enough to generate significant overlapping vocalisations, rendering it especially useful in hyperdiverse tropical ecosystems where other methods for interpreting soundscape recordings face persistent challenges.

## 1 Introduction

Estimating species richness across space and time is a foundational goal of biodiversity appraisal (Roswell et al. 2021). Biodiversity is in decline worldwide (Diaz et al. 2019), and large-scale monitoring to assess shifts in richness is critically important for understanding global change impacts (Pereira & Cooper 2006). Improving biodiversity monitoring techniques is particularly important in the tropics, which are underrepresented in ecology research (Culumber et al. 2019) despite harboring over 75% of all species (Barlow et al. 2018). The impact of environmental changes on species richness in an area can be obfuscated by simultaneous changes to the total abundance of individuals, the spatial aggregation of species (clumping), and the dominance vs evenness of individual species (Hillebrand et al. 2017). These problems can be alleviated, in part, with sampling at higher spatial and temporal resolutions (McGlinn et al. 2018). Doing this in the field is difficult, however, as the spatiotemporal resolution of traditional surveys is limited by the number of experts and parataxonomists, particularly in the tropics, each one capable of only a finite number of surveys per day (Basset et al. 2004). Bypassing the spatiotemporal constraints of traditional surveys, passive acoustic monitoring (PAM) has emerged as a leading method for biodiversity research (Sugai et al. 2019, Stowell & Sueur 2020). Early PAM research leaned heavily on manual labelling for acoustic data analyses, but this is time-consuming and limited by the availability of expert annotators. Moreover, the accessibility and reliability of analytical pipelines that do not require manual annotation remains a major barrier (Malerba et al 2026), and developing tools that can estimate species richness from recordings without manual taxonomic identification/calibration remains a priority for computational ecologists (Celis-Murillo et al. 2009, Wimmer et al. 2013).

One solution to this problem involves the use of acoustic indices, which use the spectral and temporal properties of acoustic energy in audio recordings to characterize soundscapes (Sueur & Farina 2015, Bradfer-Lawrence et al. 2019). While acoustic indices can produce valuable insights when calibrated and used correctly (Bradfer-Lawrence et al. 2025), studies have increasingly demonstrated structural issues impacting their generalisability as proxies for species richness (Sethi et al. 2023), because the relationship between variation in acoustic energy and species richness is likely too variable to be described by simple rule-based metrics (Eldridge et al. 2018, Bicudo et al. 2023, Giuliani et al. 2024, Michielsen et al. 2025, Chen et al. 2026). For example, particularly problematic in a tropical context is the reliance of many acoustic indices on the validity of the acoustic niche hypothesis (Krause 1993) which posits that acoustically active species are partitioned across time and frequency domains, and that species richness is thus a function of the diversity of acoustic information. However, tropical soundscapes often feature extensive time and frequency overlap between vocalizations of different taxa (Tobias et al. 2014, Sugai et al. 2021, Sugai et al. 2026). Additionally, the impact that individual species have on soundscape characteristics is not equal; a small number of loud and/or vocally active species can make a greater contribution to soundscape saturation than a large number of species that vocalize infrequently or in narrow frequency ranges (Diepstraten & Willie 2021).

Another approach for assessing species richness without manual annotation relies on deep learning-based animal vocalization classifiers (Stowell 2022). In particular, convolutional neural networks (CNNs) trained on thousands of bird species like BirdNET (Kahl et al. 2021) and Perch (van Merriënboer et al. 2025), have demonstrated variable ability to classify multi-species assemblages, unlocking enormous analysis potential for biodiversity research (Sethi et al. 2024, Shen et al. 2026). Despite their rapid development such models remain limited by training data availability (Freeman 2024, de Araújo 2024). Strongly annotated datasets are especially valuable (Ghani et al. 2023), and remain very rare in lowland tropical systems, with only a handful of open access examples (e.g., Hopping et al. 2022, Ruiz et al. 2026). As a result, classifiers perform much better in Europe and North America than in more species-rich Latin America, Africa, and Asia (Funosas et al. 2026, Metcalf et al. 2026). In situations where the classification performance of out-of-box models is insufficient, creating custom multi-species classifiers utilizing transfer learning is increasingly viable (Sugai et al. 2025). This is facilitated by the emergence of new tools, e.g., embedding and vector search workflows like Agile Modeling (Dumoulin et al. 2025), and advanced Bayesian occupancy models to calibrate multi-species classification outputs (Wood et al. 2024). However, despite the improved feasibility of training custom versions of these models and detectors, these remain best suited for focal taxa (Dumoulin et al. 2025) or systems with lower species richness than the humid tropics. Not only are the soundscapes in the tropics much more difficult to parse, validating classifiers for them also requires higher sampling volumes to account for the number of rare species (de Araújo et al. 2024).

Here, we develop and test a novel approach for estimating avian species richness in soundscape recordings, henceforth referred to as ‘source-clustering’, which does not 1) require manual annotation, 2) depend on variation in acoustic energy and species richness having a consistent positive relationship, like acoustic indices, or 3) need large species-specific training datasets, like classifiers. Source-clustering is designed to take advantage of the substantial advances in the feature extraction and embedding components of models like Perch and BirdNet, which remain robust across ecological systems, while circumventing the species-based training data requirements of their classifier heads, as this make them particularly vulnerable to performance declines in the tropics, where many species have low recording volumes (Kather et al. 2025). We do this by integrating cutting-edge source-separation, a machine learning approach used for e.g. determining the number of speakers in a recording (Nachmani et al. 2020), and hierarchical agglomerative clustering of the embeddings generated by Perch 2.0 from the resulting separated channels. We focus on birds, as they are the original targets of these models, but our method has potential applications for other taxa. We ask two primary research questions, applied across the temporal resolutions of 15-seconds, 1-minute, 15-minutes, and 1-hour: (1) Can our novel source-clustering approach predict avian species richness in tropical soundscapes and generalize between two separate datasets? And (2) How does source-clustering compare to both traditional acoustic indices and the off-the-shelf version of Perch 2.0, a state-of-the-art, pre-trained, multi-species bioacoustic classification model in predicting species richness?

## 2 Materials and Methods

### 2.1 Ground truth datasets

To test the viability and generalisability of source-clustering, we employed ground truth datasets from two biogeographically different Amazonian regions and habitats: *várzea* floodplain forest in Peru (henceforth PER), and non-flooded *terra firme* forest in Brazil (henceforth STM). Deployments were conducted at different times of year and using different hardware (SOM 1). For all datasets, a set of recordings from the morning hours was manually annotated by an expert observer, using Raven Pro Sound Analysis

Software (version 1.6; K. Lisa Yang Center for Conservation Bioacoustics, Cornell Lab of Ornithology, Cornell University, Ithaca, NY, USA). PER and STM_h feature “hard labels”, with bounding boxes drawn around individual vocalizations of each species, while STM_s employs “soft labels”, i.e., a list of the species present in a given recording. “STM_s_LTR” refers to low temporal resolution (15-minute) recordings, while “STM_s_HTR” refers to high temporal resolution (15-second) recordings. For more information regarding deployments, including recorder settings, see Hopping et al. (2024) for PER, Metcalf et al. (2021) for STM_s, and Metcalf et al. (2026) for STM_h.

When exact species identity was uncertain, non-species labels such as “DOSP” (Dove species) in PER, or “Ara Sp.” (*Ara* species, a genus of Macaws) in STM, were used. We used a hierarchical approach to derive species richness from individual segments up to the dataset level. For example, “DOSP” only counted as an additional species if no other dove species is recorded in a given segment, and so on.

### 2.2 Sound file processing

#### 2.2.1 Preprocessing

First, we split the recordings from STM_h and PER into 15-second intervals, to match the length of the 15-second STM_s recordings. Next, we applied a band-pass filter to isolate the 500–4000 Hz frequency band dominated by Amazonian birds (Metcalf et al. 2020, Mueller et al. 2025). Only a small minority of avian vocalizations occurred outside of this range (<10%), and an acceptable tradeoff to remove most non-avian biophony (primarily insects). However, we still performed all analysis for the full frequency range as well.

#### 2.2.2 Source-separation and embeddings

We performed source-separation by employing BirdMixIT (Fig. 1), a model trained on environmental recordings of bird calls with mixture invariant training (“MixIT”: Wisdom et al. 2020). This improves classification performance in recordings featuring overlapping vocalizations by separating sounds coming from different sources into either four or eight channels, depending on the model (Denton et al. 2022). We chose the 8-channel version because many of our 15-second files contained more than four species (4053/17751, 22.8%), but only 102 (0.57%) contained more than eight. To eliminate channels unlikely to contain birds, we applied a simple noise removal filter.

**Figure 1:**
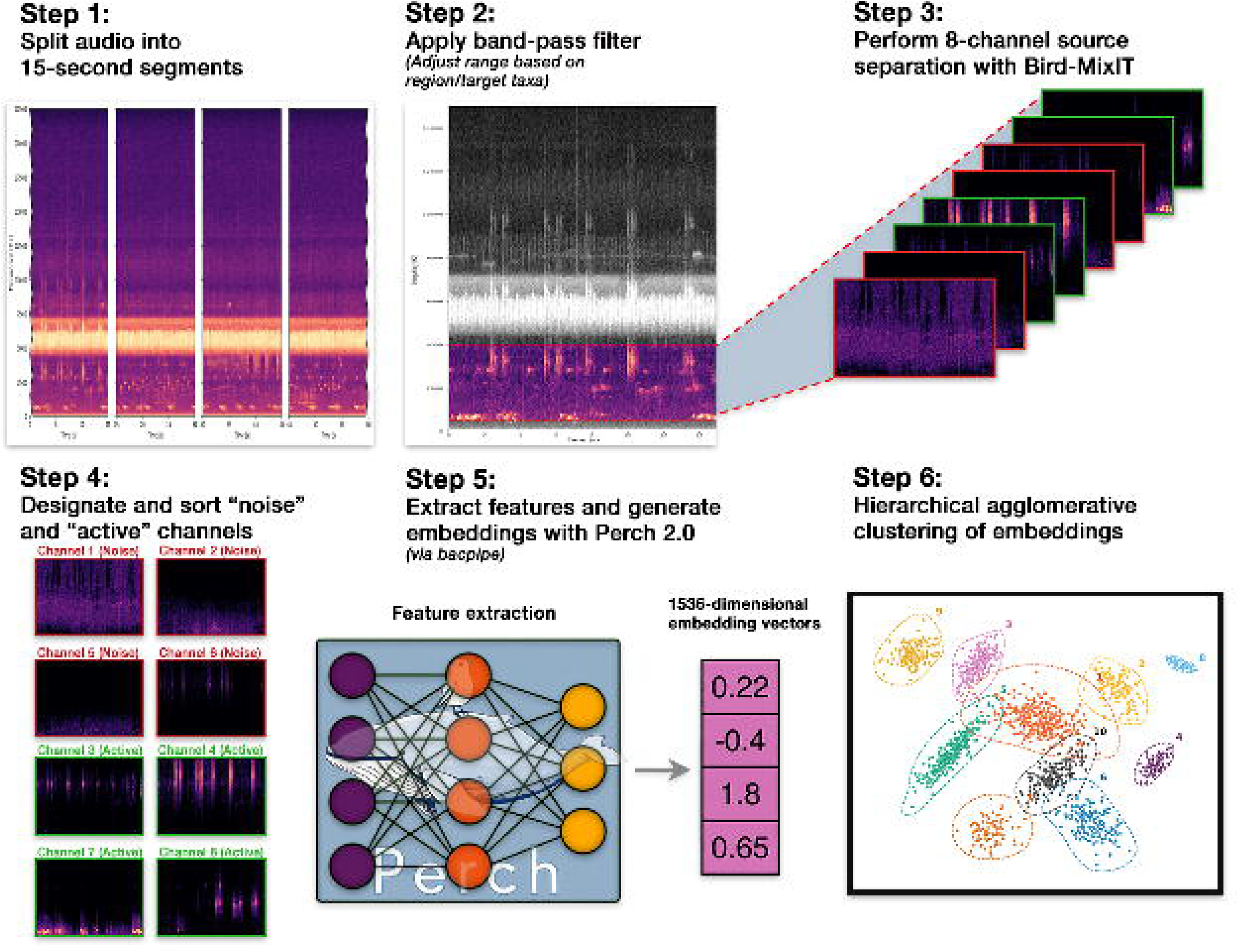
A schematic diagram of the source-clustering approach introduced in this study.

Separated channels were classed as noise if they were 1) silent or near-silent; 2) contained broadband environmental noise (e.g. rain), with very high spectral entropy and flatness; and 3) tonal insects, i.e., one narrow band dominating the channel and sustained for the entire length of it. Remaining channels were classed as “active”, and were used in our analysis. Analyses without noise filters are available in supplementals.

We chose Perch 2.0 because it performs best amongst leading bioacoustics foundation models on BirdSet (Rauch et al. 2025), a benchmark that includes our PER dataset (Schwinger et al. 2026). We did not test BirdNet because of data contamination; the most recent iteration is trained on PER. We generated feature embeddings with Perch 2.0 (van Merriënboer et al. 2025) for all separated channels using the Python package bacpipe (Kather et al. 2026). To estimate the number of different vocalization types (a species richness proxy), we sorted these embeddings into clusters using the Scikit-learn package in Python (v1.7.2, Pedregosa et al. 2011). We chose agglomerative hierarchical clustering because it was most compatible with the biological nature of the bird vocalizations in our dataset (Legendre & Legendre 2012, Chapter 8). Based on hyperparameter optimization, we calculated pairwise Euclidean distances between the *L^2^*-normalized embedding vectors, applying Ward’s minimum variance method (Ward 1963) for linkage to minimize within-cluster variance and produce compact, distinctly partitioned groupings. We defined a constant fixed distance threshold of 0.7 to terminate cluster merging.

To account for structural differences between recorder deployments, clustering was performed on embeddings pooled at the site level for the STM datasets and the site-date level for PER; one hour of recordings in both cases. We then tallied the number of unique clusters for files at the 15-second, 1-minute, and 15-minute temporal resolutions, along with the “macro” (∼1 hour) level at which they were mapped.

#### 2.2.3 Classification of individual species

We assessed the predictive performance of 10 different settings combinations of Perch 1.0 and Perch 2.0, where ER (estimated richness) was the number of detected species above a given confidence threshold (>0.1, >0.3, >0.5, >0.7, >0.9), with and without using a regional species list to filter out-of-range species. While the raw logits output by Perch 1.0 can be reliably mapped to a 0-to-1 scale using a sigmoid activation function, Perch 2.0 is trained with a softmax head that requires the probabilities of all ∼15,000 classes to sum up to a value of 1. This results in a considerable penalty when multiple birds that would otherwise have high confidence scores are vocalizing in the same segment, so out-of-box Perch 2.0 scores are less intuitive to interpret and not directly comparable with Perch 1.0. To remedy this, we calibrated our interpretation of Perch 2.0 outputs using an empirical offset value and temperature scaling parameter based on the distribution of confidence scores from Perch 1.0 (Guo et al. 2017). While Platt scaling is possible with our ground-truth data, we chose not to do it because we did not want our workflow to require manual labels.

For the regional species lists that we used to filter Perch outputs, we used the eBird checklist for the municipality of Belterra, Pará, Brazil (https://ebird.org/region/BR-PA-020/bird-list) for STM, which includes the Floresta Nacional do Tapajós and either contains or is adjacent to the STM survey sites, and the eBird checklist for Los Amigos Biological Station (CICRA, https://ebird.org/hotspot/L492606/bird-list), a well-inventoried lowland reserve in Madre de Dios, Peru, for PER. We did not use the regional list for the Department of Madre de Dios because it contains a large number of foothill and montane species that are not present in the lowlands. To avoid unrealistic precision, we chose not to use the reserve list for ITRA itself or the formal inventory list from the Santarém region (Lees et al. 2013), as such rigorous survey data is unavailable for most tropical locations.

#### 2.2.4 Traditional acoustic indices

To compare traditional acoustic indices, we calculated a range of eight commonly-used indices (Bradfer-Lawrence et al. 2019, Haupert et al. 2025) for all temporal dimensions and datasets, both with and without the band pass filter, usingscikit-maad (v1.5.1, Ulloa et al. 2021): Acoustic Complexity Index (ACI), Bioacoustic Index (BI), Surface Roughness, Spectral SNR, Spectral Events (number of events per second, “EVNspCount_sum”; fraction of events’ duration over total duration, “EVNspFract_avg”), average Spectral Activity (“ACTspfract_avg”), and Region of Interest cover (“ROIcover”). Because only PER features contiguous 1-hour recordings, and to avoid stitching together recordings from different days and/or times, we calculated the “Macro” (1 hour) level in three ways: by averaging together index values across 15-second files (STM_h, STM_s_HTR, PER); averaging across 1-minute files (STM_h, PER), and across 15-minute files (STM_s_LTR, PER).

### 2.3 Statistical analysis

We conducted statistical analysis in R (R Core Team, version 4.4.2, 2024) and made figures with ggplot2 (Wickham 2016). To assess source-clustering’s performance for predicting species richness at variable temporal resolutions and generalisability across regions (question 1), we calculated the rank order correlation (Spearman’s ρ) and p-values for the relationship between the number of clusters produced by source-clustering (estimated richness) and observed richness derived from our manual annotations. We assessed this relationship for both PER and STM at the 15-second, 1-minute, 15-minute, and “macro” (1-hour) resolutions. For all results subsections, the 15-second resolution features PER, STM_h, and STM_s_HTR; the 1-minute resolution features PER and STM_h; the 15-minute resolution features PER and STM_s_LTR, and the “macro” (1-hour) resolution uses the site-date level for PER and the site level for STM_h, STM_s_HTR, and STM_s_LTR as independent datapoints, all of which are approximately or exactly one hour.

To evaluate the comparative performance of source-clustering, traditional acoustic indices, and various settings of the Perch 2.0 classifier for estimating species richness across the four assessed temporal resolutions (question 2), we employed a weighted bootstrapping procedure. A bootstrap sample of equal size to the full dataset at a given temporal resolution was drawn with replacement for 500 iterations, after which we fit a simple linear regression model using the acoustic metric to predict the observed species richness (SR). We then calculated the Root Mean Square Error (RMSE), weighted for effort in minutes, where absolute error was defined based on the residuals of our model (the difference between richness estimated by the model and observed SR). To standardize our comparisons across different temporal scales, we converted RMSE to relative RMSE (rRMSE) by dividing the absolute RMSE by the average observed SR of each bootstrap sample. We assessed the predictive performance of each metric by deriving and plotting the 95% confidence intervals for these rRMSE values across the bootstrap samples.

## 3 Results

### 3.1 Source-clustering and species richness

We found that source-clustering effectively predicted avian species richness at all four temporal resolutions (15-seconds, 1-minute, 15-minute, and “macro” (1-hour)) and generalized well between both datasets (Fig. 2). Based on Spearman’s ρ, dataset-specific performance ranged from a low of ρ = 0.351 (PER, 15-second resolution, p < 0.001) to a high of ρ = 0.843 (STM, “macro” (1-hour) resolution, p < 0.001). Excluding the macro level where the sample size for PER was too small for significance (n = 21, p = 0.115), we did not find that clustering performed substantially better for one dataset than the other; performance was better for STM at 15-seconds and 15-minutes, while PER saw better performance at 1-minute resolution. In general, predictive performance improved at larger temporal intervals; with STM and PER treated as a single combined dataset, ρ = 0.465 at 15-seconds, 0.525 at 1-minute, 0.646 at 15-minutes, and 0.793 at 1-hour (p < 0.001 at all levels). While STM_h had higher total species richness than PER, primarily because of the larger number of sites (and therefore higher gamma diversity), PER had higher alpha diversity and higher annotation density, averaging 14.73 and 17.16 labels/minute with and without background vocalizations, respectively, compared to STM_h’s value of 10.73 labels/minute.

**Figure 2:**
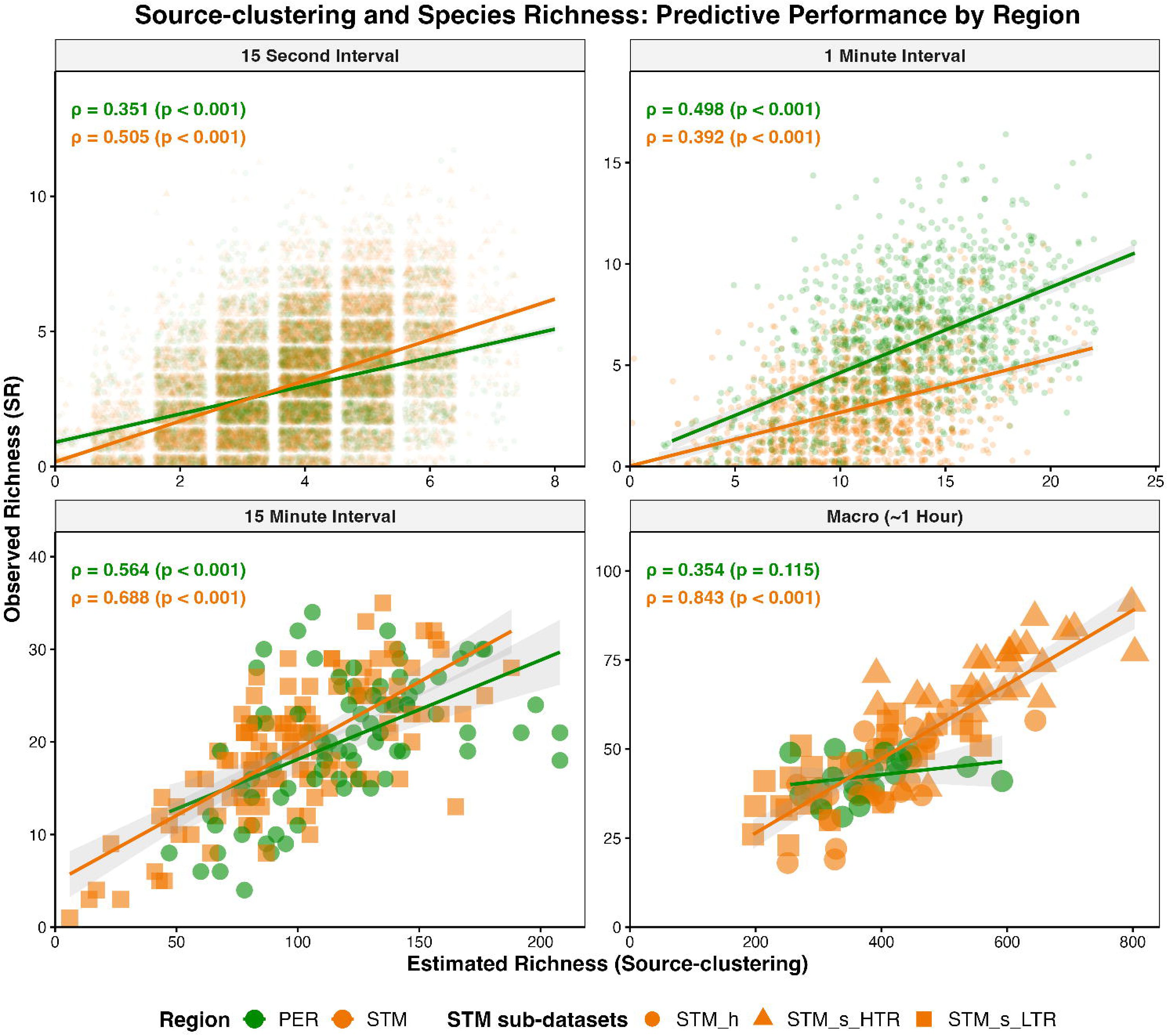
The relationship (Spearman’s ρ) between manual-annotation-derived (observed) species richness and species richness estimated by source-clustering (euclidean distance, 0.7 fixed threshold) at four temporal resolutions for two different datasets. Data shown concerns embeddings of source-separated files with the 500-4000Hz band-pass filter applied and “noise” channels removed. See Table 1 for sample sizes at each temporal resolution. Trendlines are derived from a linear model (SR ∼ ER) for each dataset.

### 3.2 Comparing source-clustering with out-of-box Perch and traditional acoustic indices

Source-clustering predicted avian species richness better than every assessed acoustic index and most settings of Perch, though it was generally comparable with the latter (Fig. 3). The gap in performance between source-clustering and traditional indices was greatest at longer temporal windows; at the 15-minute and 1-hour resolutions. Even without a band-pass filter or removal of noise channels, source-clustering still predicted species richness better than all acoustic indices, including those that incorporated their own band-pass filter.

**Figure 3:**
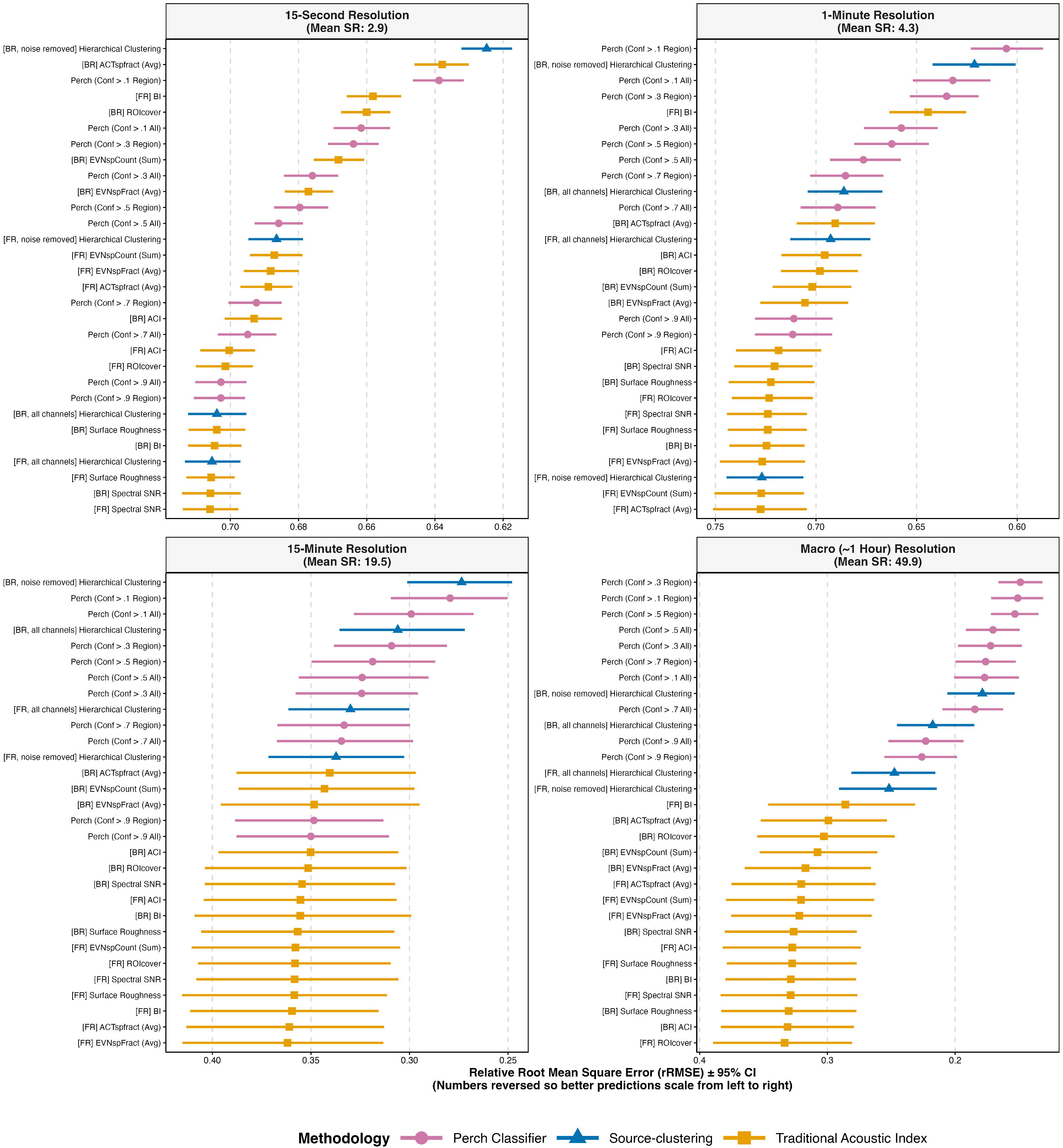
Predictive power of different Perch 2.0 settings, traditional acoustic indices, and source-clustering at 15-second, 1-minute, 15-minute, and 1-hour temporal resolutions. rRMSE ± 95% CIs were based on 500 bootstrapping iterations of the difference between observed species richness and species richness estimated based on a simple linear regression model for each metric, weighted for effort in minutes, and relativised by dividing raw RMSE values by the average species richness of the bootstrap sample. Estimated richness from Perch 2.0 is calculated based on the number of detections above each adjusted confidence threshold, with and without a regional species filter. Metrics are labeled [BR], “bird range”, when a band-pass filter isolating the 500-4000Hz frequency range is applied, and [FR], “full range”, without band-passing. Featured acoustic indices are defined in Section 2.2.4.

In general, source-clustering performed better than Perch 2.0 with the most liberal out-of-box settings applied (confidence >0.1, no regional species list, Fig. 4). For other settings and comparison with Perch 1.0, see SOM 2. Perch 2.0 almost always performed better than Perch 1.0, though the gap between them shrunk considerably at higher confidence thresholds; the difference in average R^2^ between Perch 1.0 and Perch 2.0 was 0.084 at >0.1, 0.034 at >0.3, 0.0366 at >0.5, 0.024 at >0.7, and just 0.014 at >0.9. Source-clustering performed better than Perch 2.0 in all metrics (R^2^, RMSE, MAE) at the 15-second and 15-minute resolutions, and at 1-minute, source-clustering performed better than all settings of Perch 2.0 other than confidence >0.1 with a regional species list applied. At the macro (∼1 hour) level, most settings of Perch 2.0 performed better than source-clustering, though without a regional species list, Perch 2.0 performed only slightly better than source-clustering at confidence intervals of >0.1, >0.3, and >0.5, and worse at >0.7 and >0.9. Overall, Perch, like source-clustering, saw its predictive performance improve generally at longer temporal intervals, but the decline in performance it experienced at narrow temporal resolutions was steeper than clustering. Additionally, lower Perch confidence thresholds consistently produced stronger predictive values, though >0.3 performed better than >0.1 for both Perch settings at the “macro” (1-hour) level.

**Figure 4:**
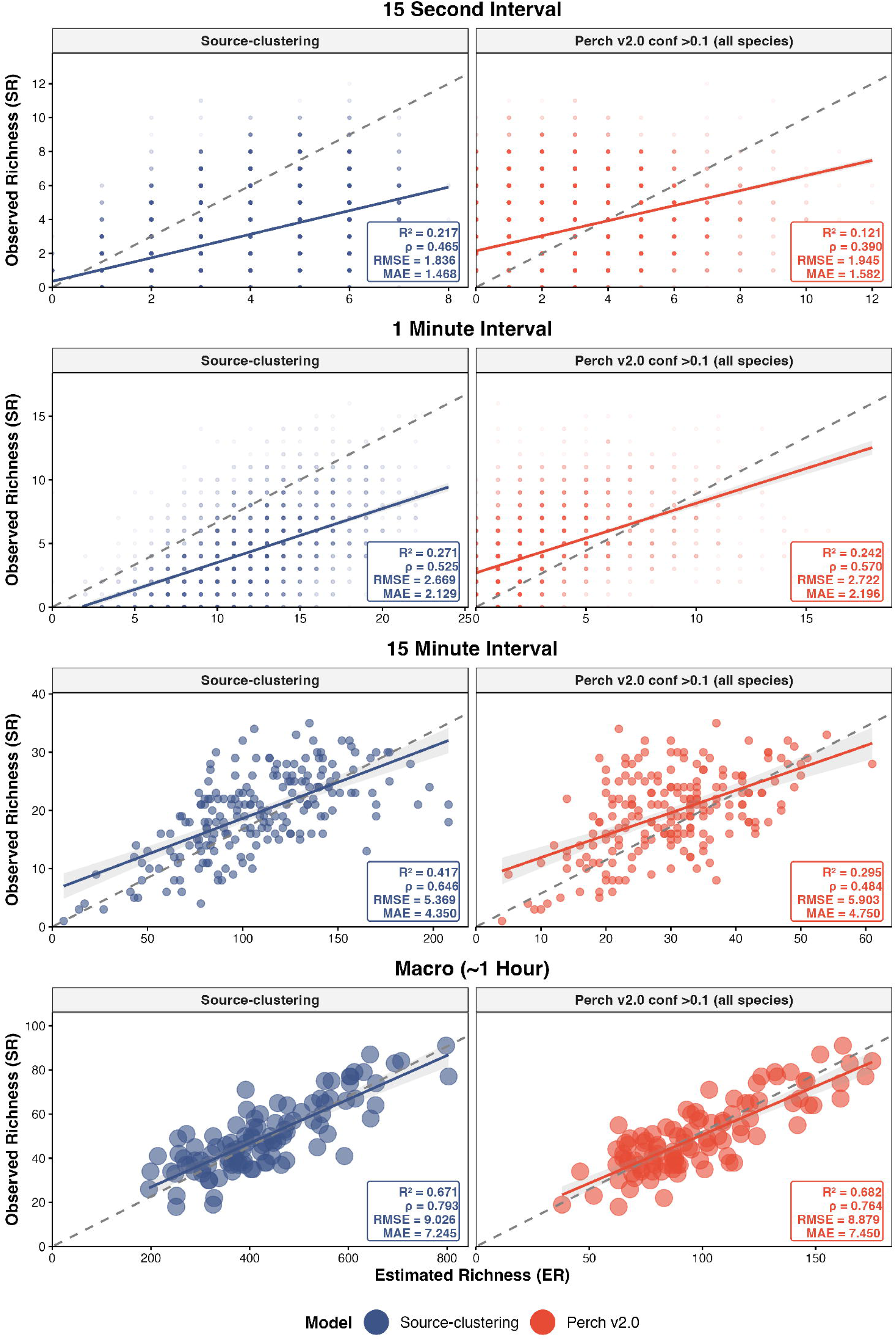
Comparison of observed (annotated) vs estimated species richness, via source-clustering and classifier outputs from Perch 2.0, at a variety of temporal resolutions. Species richness estimated by Perch is the sum of detected species with an adjusted confidence score of >0.1, without a regional list applied.

## 4 Discussion

### 4.1 Applications and advantages of source-clustering

We believe that the value of source-clustering is likely greatest for analysing recordings at high temporal resolutions and in remote and/or hyperdiverse systems, particularly the tropics, where classification models tend to struggle because of the large amount of overlapping vocalizations and insufficient availability of training data. That the marginal advantage of source-clustering was greatest for shorter recordings contributes to its value in such systems, because PAM arrays in remote areas are often programmed to record in shorter intervals as a way of maximising deployment length.

By using a source-separation model trained specifically on bird vocalizations rather than capturing general acoustic properties of recordings, this approach retains one of the key advantages that species-based soundscape analysis holds over traditional acoustic indices. At the same time, it complements species-based approaches, as it does not require individual species classification. There are many global systems where training data volume, and therefore expected classifier performance, is considerably lower than the Amazon, including most of tropical Africa and Asia (Funosas et al. 2026). In these settings, our clustering method is likely to perform better than classifiers at all temporal resolutions, as its underlying architecture should be much less sensitive to shifts between regions than classifiers that rely on the amount of training data they have for each species. Source-clustering can even replace the need for manual annotation in some cases. For example, fig. 2 shows that the conclusions of Metcalf et al. (2021) could have been reached without requiring manual annotation at all, by applying source-clustering on the same dataset; both the cluster count and species richness values of STM_s_LTR segments are consistently lower than those of STM_s_HTR at the 1-hour resolution, which was the primary finding of that study.

### 4.2 Reflections on classifier outputs

When comparing our method to classification results, we primarily focused on Perch without a regional species list and with a confidence filter of >0.1 to get as many detections as possible. Classifier performance for predicting species richness in Amazonia has been found to be strongest with lower confidence intervals (Pérez Granados et al. 2025, Metcalf et al. 2026), though this could change as classifiers improve, and ultimately we only knew that low confidence thresholds were best for estimating species richness in our dataset because we had extensive manual annotations to test against, which is not the case for most PAM studies. We felt that aggregating detections without a regional species list was a more appropriate comparison with the clustering workflow, which does not use species-level identification at all, and regional species lists are neither an out-of-box Perch feature nor available at consistent quality in many tropical regions. Against these “realistic” versions of Perch, source-clustering performed especially well.

That Perch performed particularly poorly for estimating species richness at the highest confidence threshold (>0.9) is notable because some recent papers that use classifier outputs as the independent variable, i.e., as species richness, for evaluating other metrics employ the same high >0.9 confidence threshold to reduce false positives (e.g. Hutschenreiter et al. 2025). Our results suggest that this likely diminishes the effectiveness of classifier outputs as a proxy for species richness and should therefore be avoided. Because detector output scores are not transferable among species (Wood & Kahl 2024), optimal thresholds may vary by region, season, and time of day. The low performance of classifiers for estimating species richness at high confidence thresholds (especially >0.9) was exacerbated because the improvement from Perch 1.0 to Perch 2.0 was much lower at this threshold than at >0.1, which we suspect to be a result of the simultaneous singing penalty from Perch 2.0’s softmax head. This structure makes Perch 2.0 suboptimal for datasets like ours with many overlapping vocalizations, and less intuitive than Perch 1.0 to interpret in general. We believe that our calibration of Perch 2.0 outputs was effective; it showed that Perch 2.0 improved on Perch 1.0 by roughly the same margin as estimated in Perch 2.0’s introduction paper (van Merriënboer et al. 2025).

Our comparisons with Perch use out-of-box versions rather than a custom, tuned classifier, which is theoretically optimal, but we feel that out-of-box settings present a more realistic example of how these models are likely to be used at present. To date, efforts to create custom multi-species classifiers for individual studies or projects are best suited for less species rich assemblages than lowland tropical rainforests. For instance, the examples of bespoke multi-species classifiers given in Sugai et al. (2025) concern 67 diurnal birds (Wood et al. 2024), 42 frog species (Cañas et al. 2023), 31 katydids (Madhusudhana et al. 2024), and 12 birds and 2 mammals (Ruff et al. 2021).

All of these studies, combined, form a considerably smaller species pool (136 species) than are featured in any of the three main datasets we used in this paper; PER has 170 species, STM_s has 245 species, STM_h includes 195 species, and STM and PER together have 370 (SOM 1), themselves incomplete site inventories. When they are created for the tropics, such models typically only concern a small minority of present species (Dubus et al. 2026). Creating complete multi-species classifiers in systems as complex as the Amazon would be a serious undertaking and isn’t presently realistic for the small scale and budget of many PAM studies though sophisticated new tools like vector search with Agile Modeling (Dumoulin et al. 2025) continue to make this endeavor more realistic. Similarly, sophisticated Bayesian multi-species occupancy modelling techniques can account for imperfect detection and misclassification in classifier outputs, but require manually validated predictions at the species level and consistent hardware and recording settings (Wood et al. 2024). As a result, such approaches are most applicable for large projects with uniform hardware, disproportionately located in the Global North (Sánchez Herrera et al. 2024), and in systems with fewer species; Wood et al. (2024) were only able to validate about 80 species to the full extent, in the well-studied Sierra Nevada. Doing so would be even more difficult in complex tropical assemblages, given the limited pool of experts (Robinson et al. 2018). Even fewer of these field identification experts have access to the computational resources and expertise necessary to fully leverage emerging analysis approaches, though source separation can also be computationally expensive.

### 4.3 Pathways forward

Improvement in the performance of our approach is likely to be achieved most readily with more sophisticated noise removal (e.g. Denton et al. 2022). Our noise removal approach was rudimentary in order to maximise simplicity and replicability while still outperforming traditional indices - training a model specifically for identifying noise in separated channels was outside of the scope of this study. However, a noise-identification model could be trained using semi-supervised classification of separated noise channels or by inputting all channels of recordings that are known not to contain birds. Even with basic noise removal, however, our method still performed better for estimating species richness than every assessed traditional acoustic index at all temporal resolutions, and compared equivalently or favorably to Perch, a gap that better noise detection and removal could widen substantially. While removing noise channels improved results, doing so was not necessary to achieve higher scores than most traditional acoustic indices and some Perch settings at the 1-minute, 15-minute, and 1-hour resolutions (see the “BR, all channels” values in Fig. 3), which speaks to the effectiveness of our hierarchical aggregative clustering method.

Our approach represents an intermediate step on the way to full multi-species classification in all systems, rather than a tangential output like acoustic indices tend to be. Instead of producing standalone index values, retaining embeddings for each separated segment enables more sophisticated analysis like assessments of similarity between sites, and a classification head can be applied to these embeddings. Denton et al. (2022) found that overall classification performance declined on separated channels, likely due to the absence of audio context, improving when the original mixture was included as an additional channel, but this used a classifier trained on recordings that had not undergone separation. Training a classifier specifically on audio that has been separated and had noise channels removed, like we do in this paper; integrating that process into the model for input recordings, could improve performance considerably, particularly for overlapping vocalisations.

As with noise channel removal, applying a band pass filter at the 500-4000Hz range improved performance considerably, but was not necessary for the predictive performance of clustering to exceed that of all calculated indices at the 15-minute and 1-hour temporal resolutions. As we demonstrate with its impact on predictions of overall avian species richness, an adjusted band pass filter can be used to target specific taxa that occupy defined frequency ranges, like bats. However, relying on a band pass filter to isolate birds does limit the performance ceiling of our approach by excluding species that vocalize outside of the 500-4000Hz range, though in the case of the Amazon, these vocalizations disproportionately belong to groups like tanagers and hummingbirds that are both difficult for human annotators to identify, and, as a result of being poorly known, feature low training data volume for classifiers. Likewise, while in Amazonia the 500-4000Hz frequency range is dominated by birds during the morning, a handful of insects and frogs do vocalize in this range, and distinguishing all of them from birds is not possible with source-clustering alone – doing so would require a classifier head.

Determining the frequency range for the band pass filter depends on specific knowledge of the study system, and while the Amazon presents an unusually well-defined range, this is likely a function of acoustic niche partitioning with the Amazon’s abundant insects (Metcalf et al. 2020). Systems where the frequency range for birds is not as strongly defined also tend to feature reduced interference from insects, so isolating said frequency range is therefore less important.

## Supporting information

SOM 1

SOM 2

## 5 Conclusion

We present a novel method for estimating avian species richness in soundscape recordings, using sound separation and agglomerative hierarchical clustering of embeddings to provide a useful advance on acoustic indices. Our approach, source-clustering, can complement or even outperform leading out-of-box automated species classification models in hyperdiverse tropical forests. By utilising the components of sophisticated classification models that are most effective (feature embedding) without being subject to the location-specific training data requirements that can impair their classification performance, especially in the tropics, where effective PAM analysis methods are arguably most important. Source-clustering should generalize better across remote and under-studied systems than alternative methods, with significant room for upwards improvement via enhanced noise removal, or even the development of classifiers that incorporate source-separation as part of their training or data processing pipelines.

## Acknowledgements

We thank the Large Scale Biosphere-Atmosphere Program and the Inkaterra Association for logistical and infrastructure support during field work in Brazil and Peru, respectively, as well as Joice Ferreira for her support in Brazil. We are grateful for the hard work of our field and laboratory assistants in Brazil: Gilson de Jesus Oliveira, Renílson Freitas, Josué de Jesus Oliveira, and Josiane de Jesus Oliveira; and in Peru: Noe Roger Huaraca-Charca, Dennis Osorio, and Kevin Jiménez Gonzales. We thank Fernando Angulo and Ken Rosenberg for helping to identify unknown vocalisations in the Peru dataset; Sidnei Dantas and Mario Cohn-Haft for Brazil. Lastly, we are very grateful for the engagement of Larissa Sayuri Moreira Sugai, Vincent Kather, Inês Nolasco, and Vincent Dumoulin, who we bounced ideas off of during the early conceptual stages of the paper.

## Authors’ contributions

WAH conceived of the idea of the paper and designed methodology with the significant assistance of JB, SS, OCM, and ACL; WAH (Peru) and OCM, EB, and LCR (Brazil) collected and curated the data; WAH (Peru) and NGM (Brazil) with help from ACL annotated the ground truth datasets; and WAH analysed the data and led the writing of the manuscript. All authors contributed critically to the drafts and gave final approval for publication.

## Data Availability Statement

The code used to perform this analysis, along with a spreadsheet including the raw data values for observed species richness, cluster counts, computed acoustic indices, and Perch-derived estimated species richness of every recording interval analyzed were uploaded as part of the peer review process, and can be made public upon acceptance. Additionally, the full recordings and annotations composing the PER dataset are fully open-sourced: see zenodo.org/records/7079124.

