## Supplementary material for "Unsupervised source separation and clustering of soundscape recordings predicts avian richness in hyperdiverse systems": SOM 1

**Table 1.** Recorder deployment details for each dataset.

| Dataset | Location | Label type | Recorder type | Citation | Dates | Time of day |
| --- | --- | --- | --- | --- | --- | --- |
| PER | Inkaterra Reserva Amazónica (ITRA)<br>(12°32′07.8″ S, 69°02′58.2″ W),<br>Madre de Dios, Peru | Hard labels | Swift Recorder (1st<br>generation) | K. Lisa Yang Center for<br>Conservation Bioacoustics,<br>Cornell Lab of Ornithology,<br>Ithaca, NY, USA | 14 January to 2<br>February 2019 | 05:00–06:00 (dawn) |
| STM (all) | Municipalities of Santarém, Belterra,<br>and Mojui dos Campos (~3°02′45.6″<br>S, 54°56′49.2″ W), Pará, Brazil | Mixed | Mixed | – | – | Spread across 3 hours<br>starting 30 minutes<br>before dawn |
| STM_h | *See STM (all)* | Hard labels | AudioMoth v1.2 | Hill et al., 2018 | 22 November to 14<br>December 2022 | *See STM (all)* |
| STM_s | *See STM (all)* | Soft labels | Frontier Labs<br>Bioacoustic Recording<br>Units | Frontier Labs, 2015 | 12 June to 16 August<br>2018 | *See STM (all)* |
| STM_s_LTR | *See STM (all)* | Soft labels | Frontier Labs<br>Bioacoustic Recording<br>Units | Frontier Labs, 2015 | 12 June to 16 August<br>2018 | *See STM (all)* |
| STM_s_HTR | *See STM (all)* | Soft labels | Frontier Labs<br>Bioacoustic Recording<br>Units | Frontier Labs, 2015 | 12 June to 16 August<br>2018 | *See STM (all)* |

**Table 2.** Species, site, label, and recording-effort summary for each dataset.

| Dataset | Species total | Site count | Total labels/IDs | Total recording length (h) | Number of segments |  |  |  |
| --- | --- | --- | --- | --- | --- | --- | --- | --- |
|  |  |  |  |  | 15-second | 1-minute | 15-minute | 1-hour |
| PER | 165 | 7 | 18,559 | 21 | 5,040 | 1,260 | 84 | 21 (continuous recordings) |
| STM (all) | 277 | 30 | – | 82 | 12,711 | 1,440 | 116 | 82 (aggregated at the site/dataset level) |
| STM_h | 195 | 24 | 15,450 | 24 | 5,760 | 1,440 | N/A | 24 (aggregated at the site level) |
| STM_s | 245 | 29 | – | 58 | N/A | N/A | N/A | 58 (aggregated at the site/dataset level) |
| STM_s_LTR | 204 | 29 | 2,217 (cumulative IDs) | 29 | N/A | N/A | 116 | 29 (aggregated at the site level) |
| STM_s_HTR | 224 | 29 | 27,097 (cumulative IDs) | 29 | 6,951 | N/A | N/A | 29 (aggregated at the site level) |
| **Total** | – | **37** | **34,009 (hard)/29,314 (soft)** | **103** | **17,751** | **2,700** | **200** | **103** |

Dashes (–) indicate values not reported for that dataset.
