## Supplementary figures and images for "Unsupervised source separation and clustering of soundscape recordings predicts avian richness in hyperdiverse systems"

### SOM 2

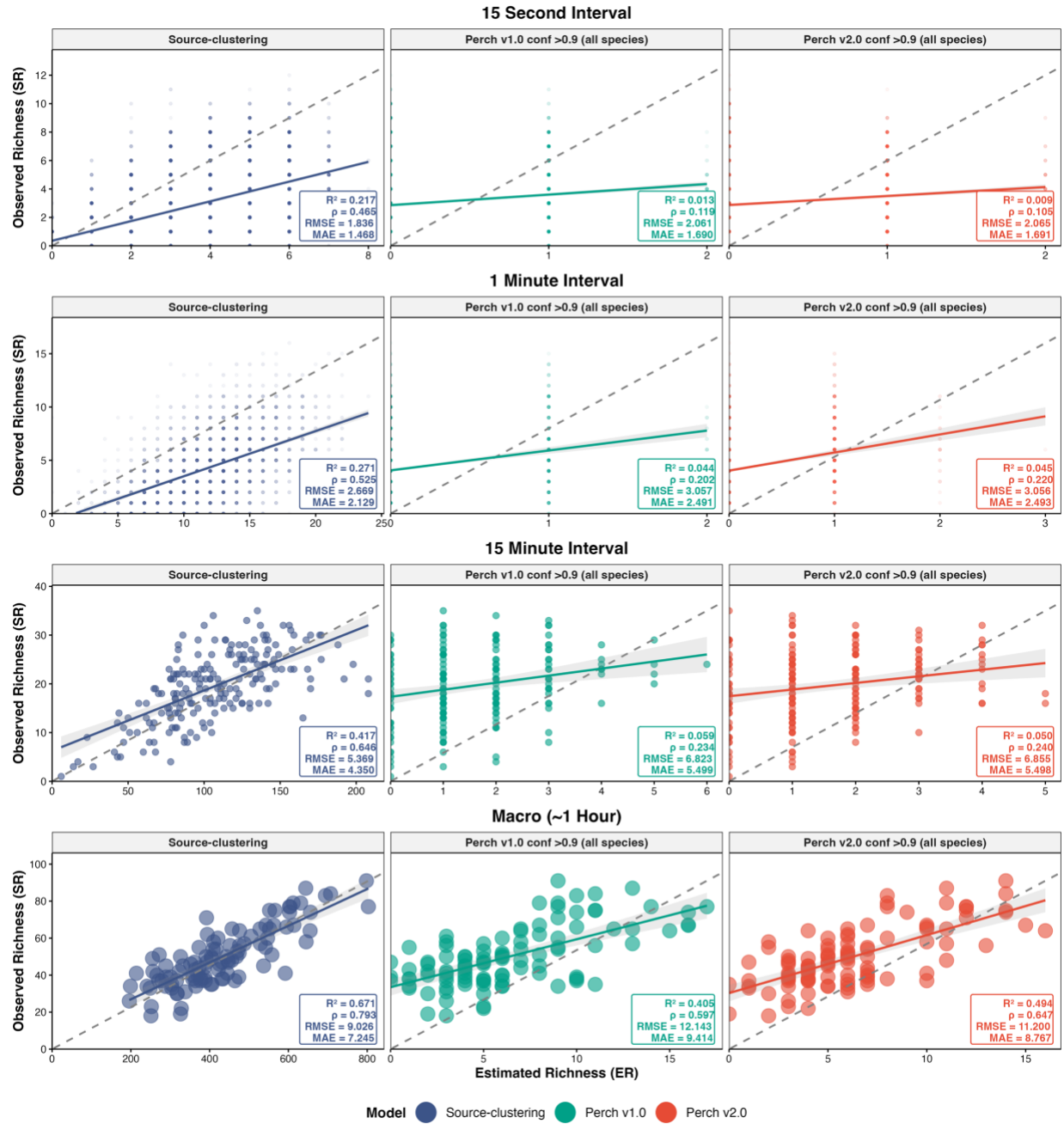

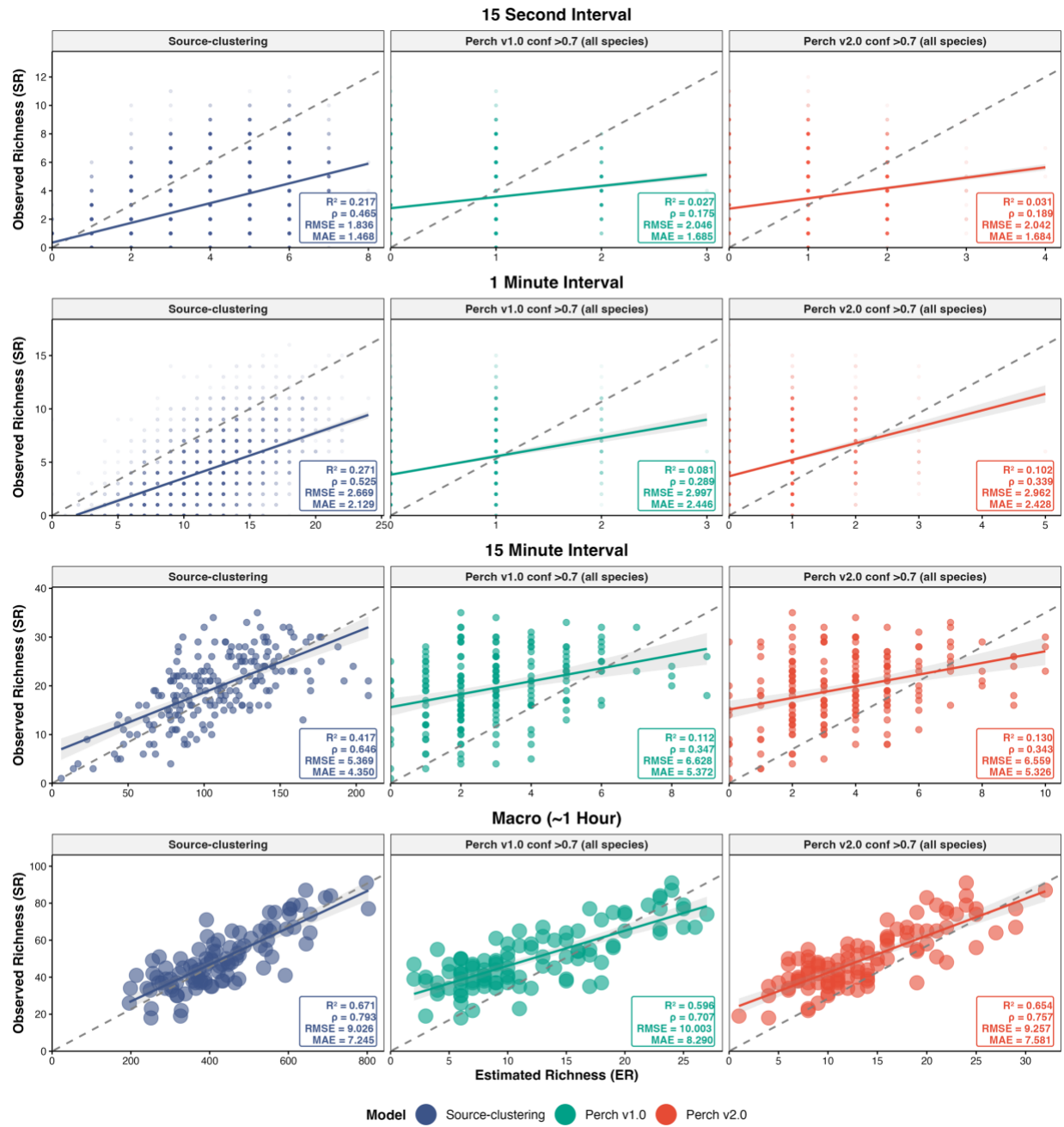

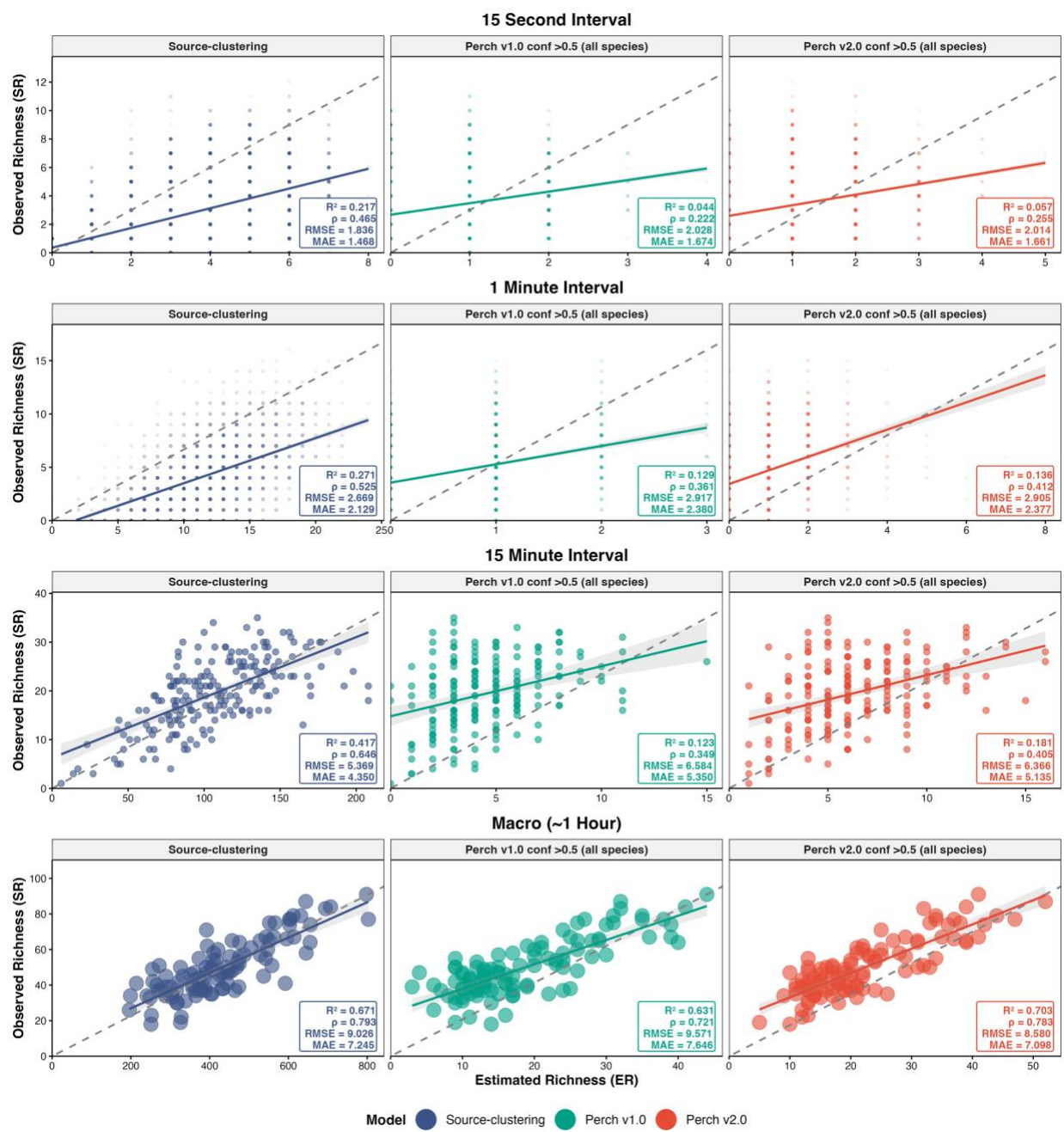

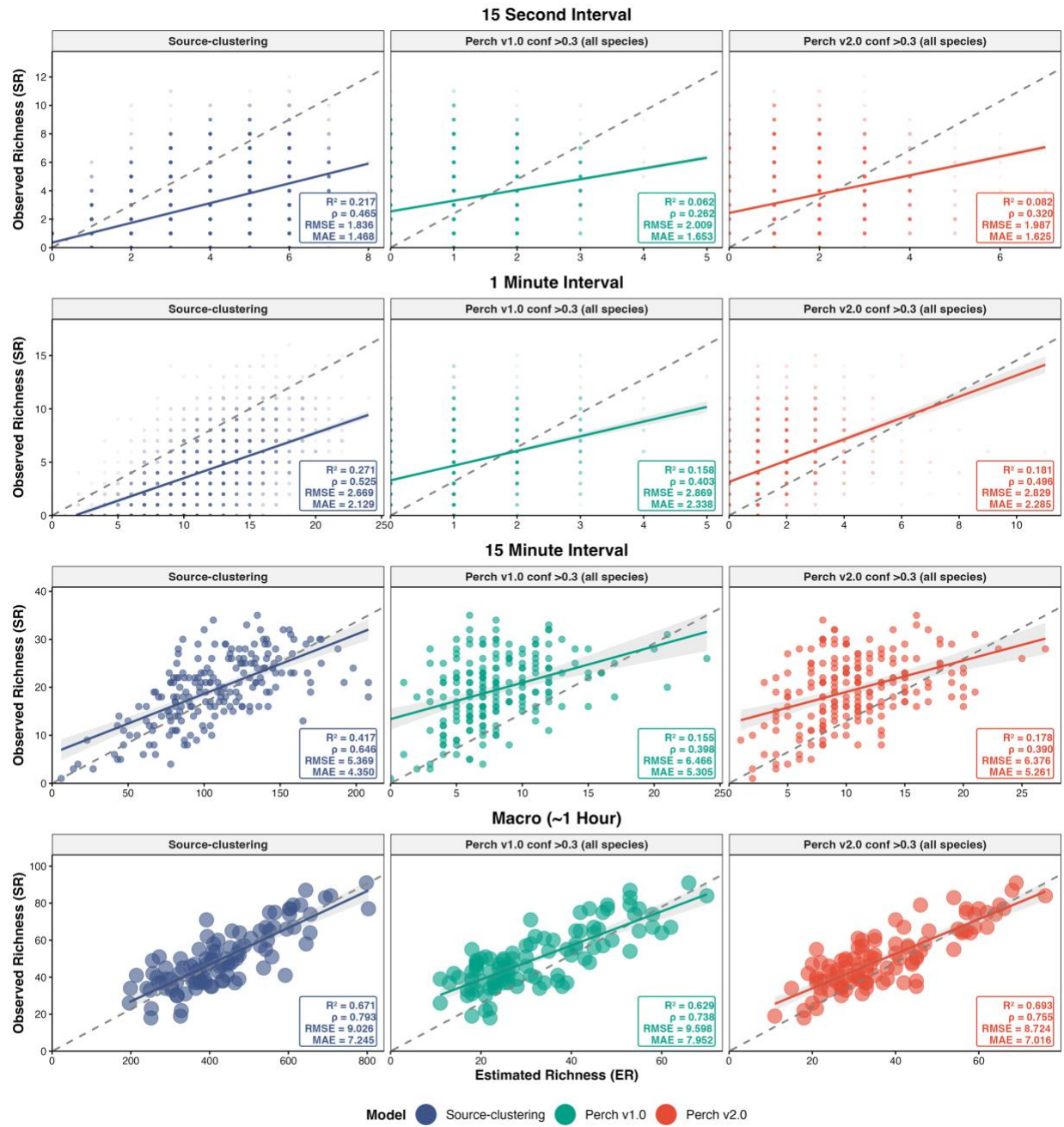

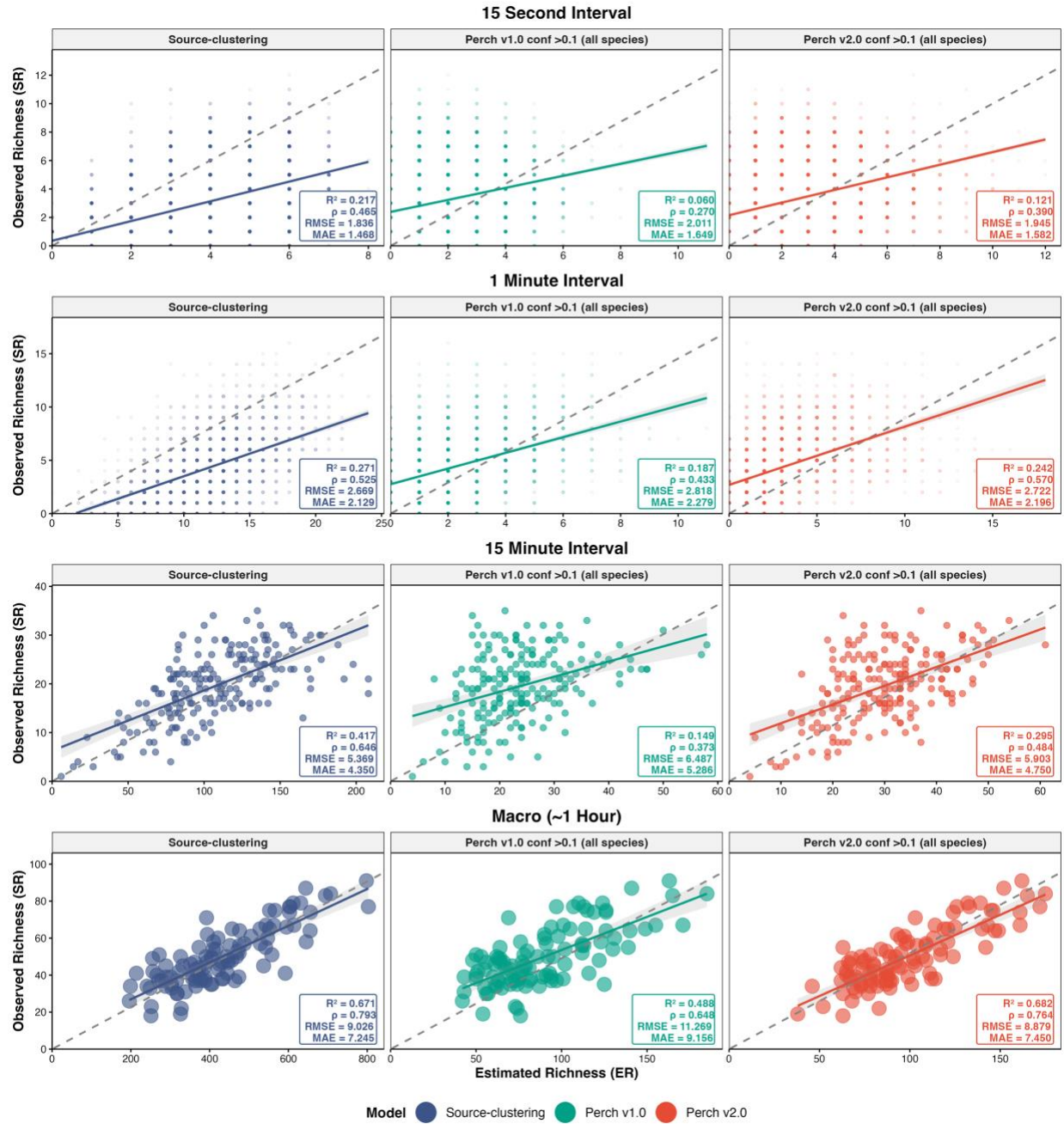

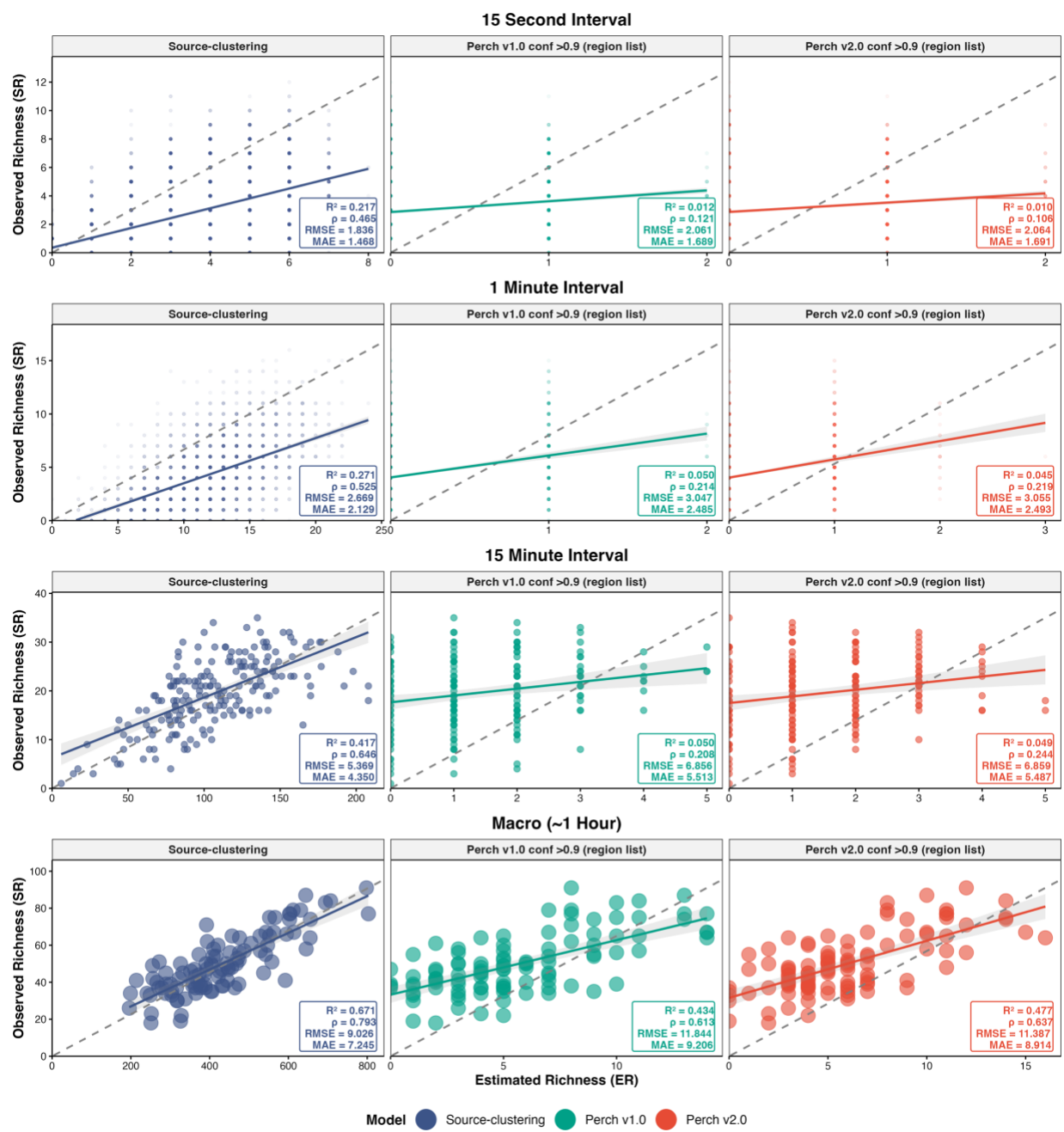

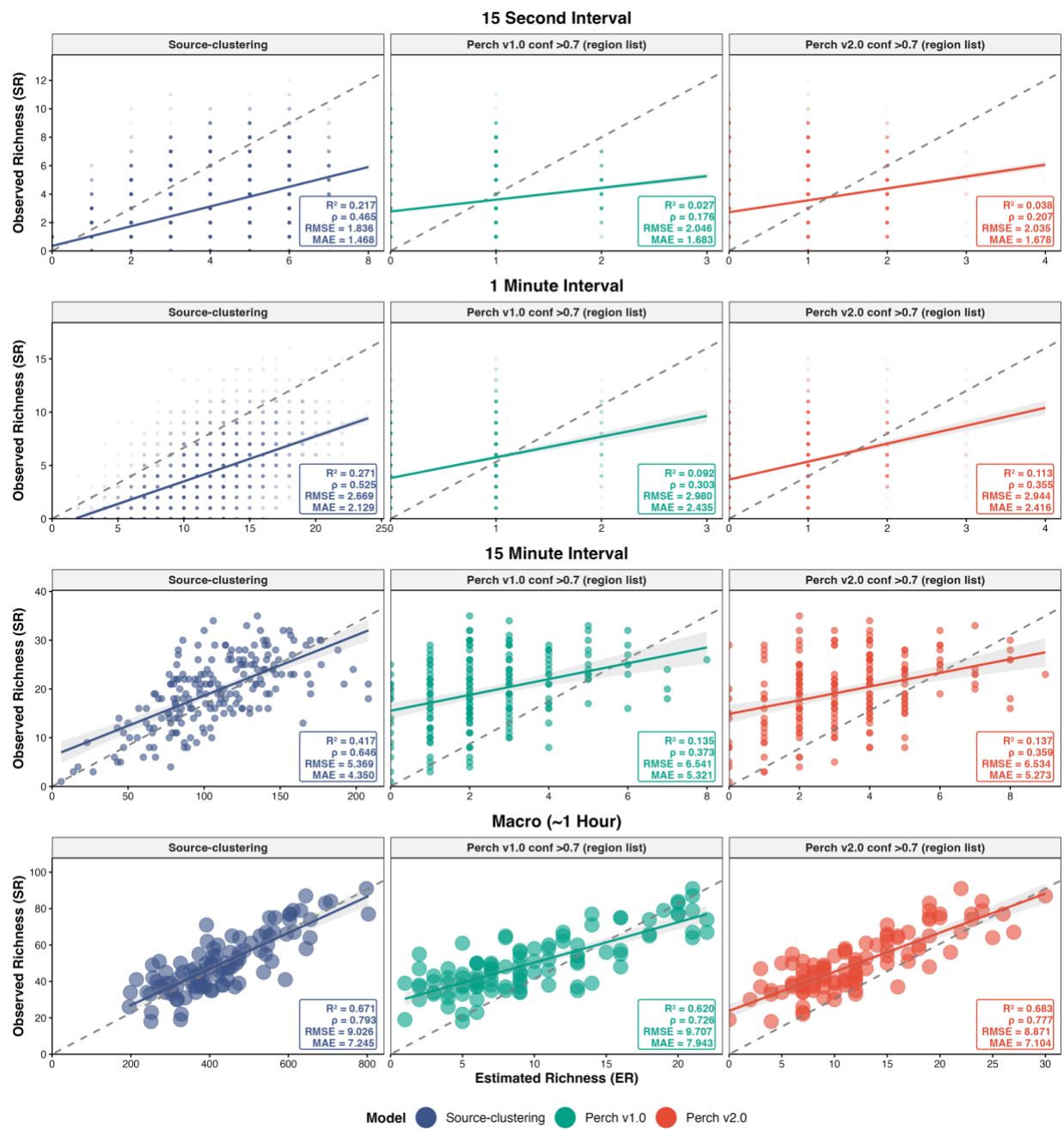

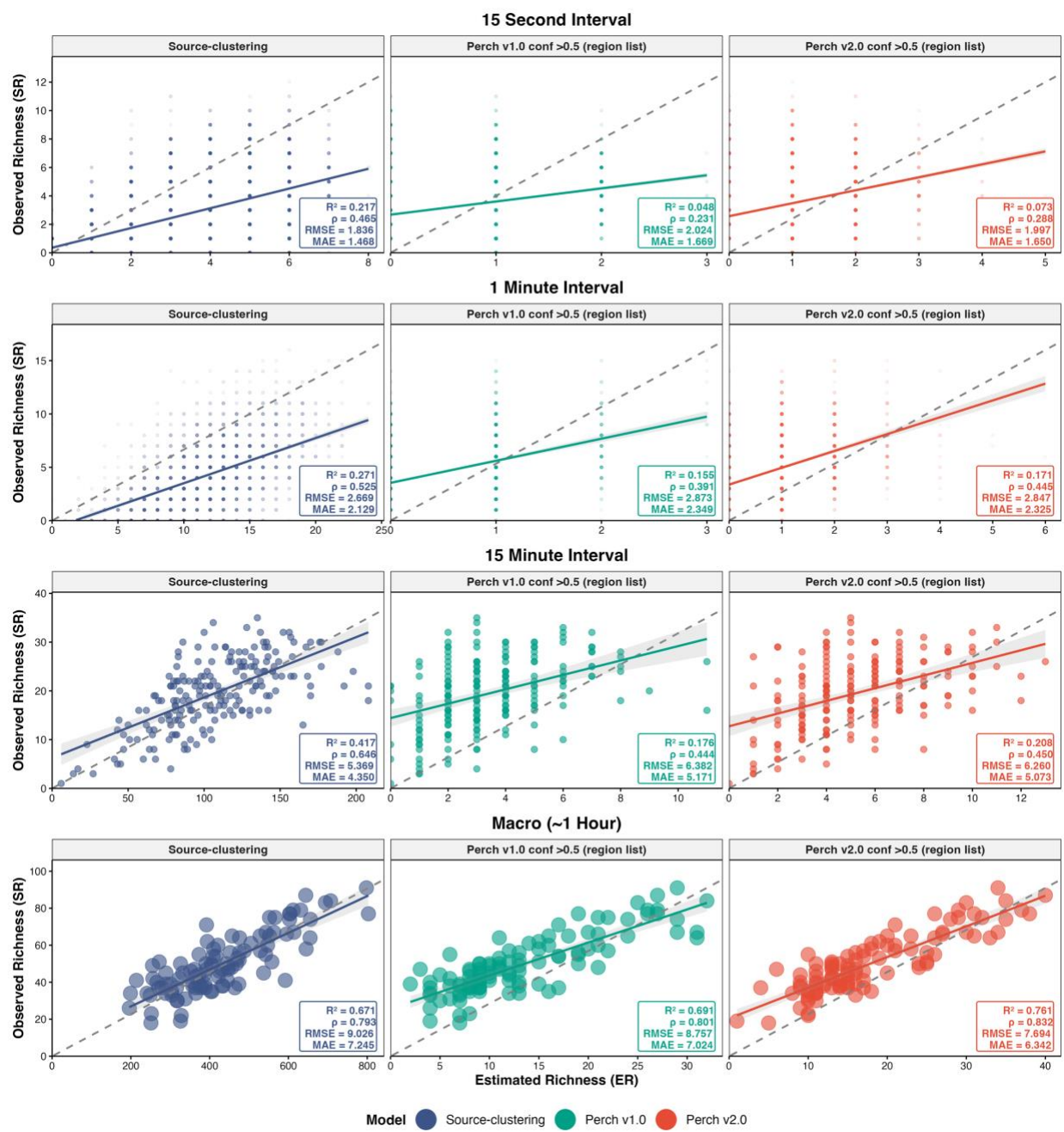

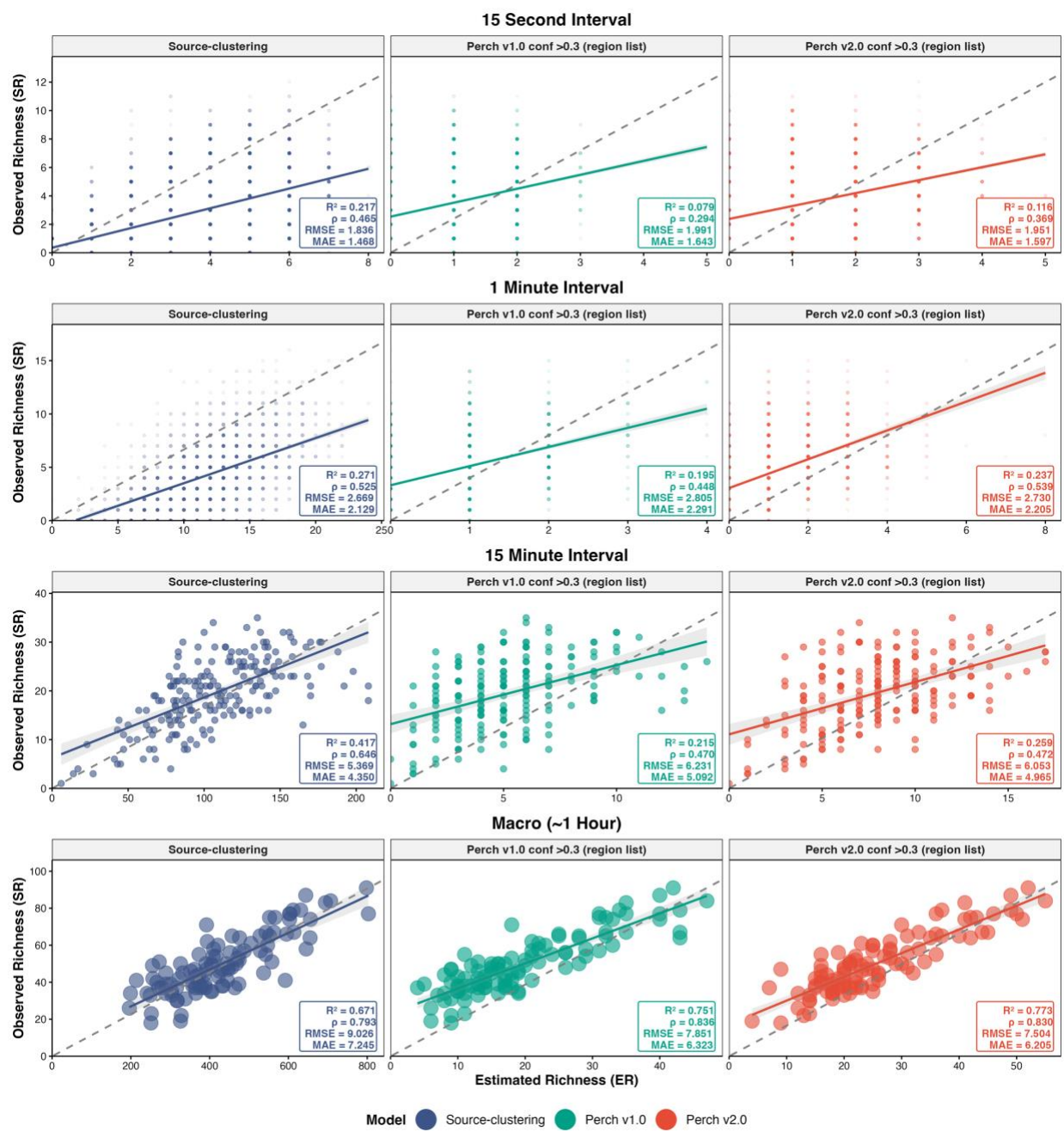

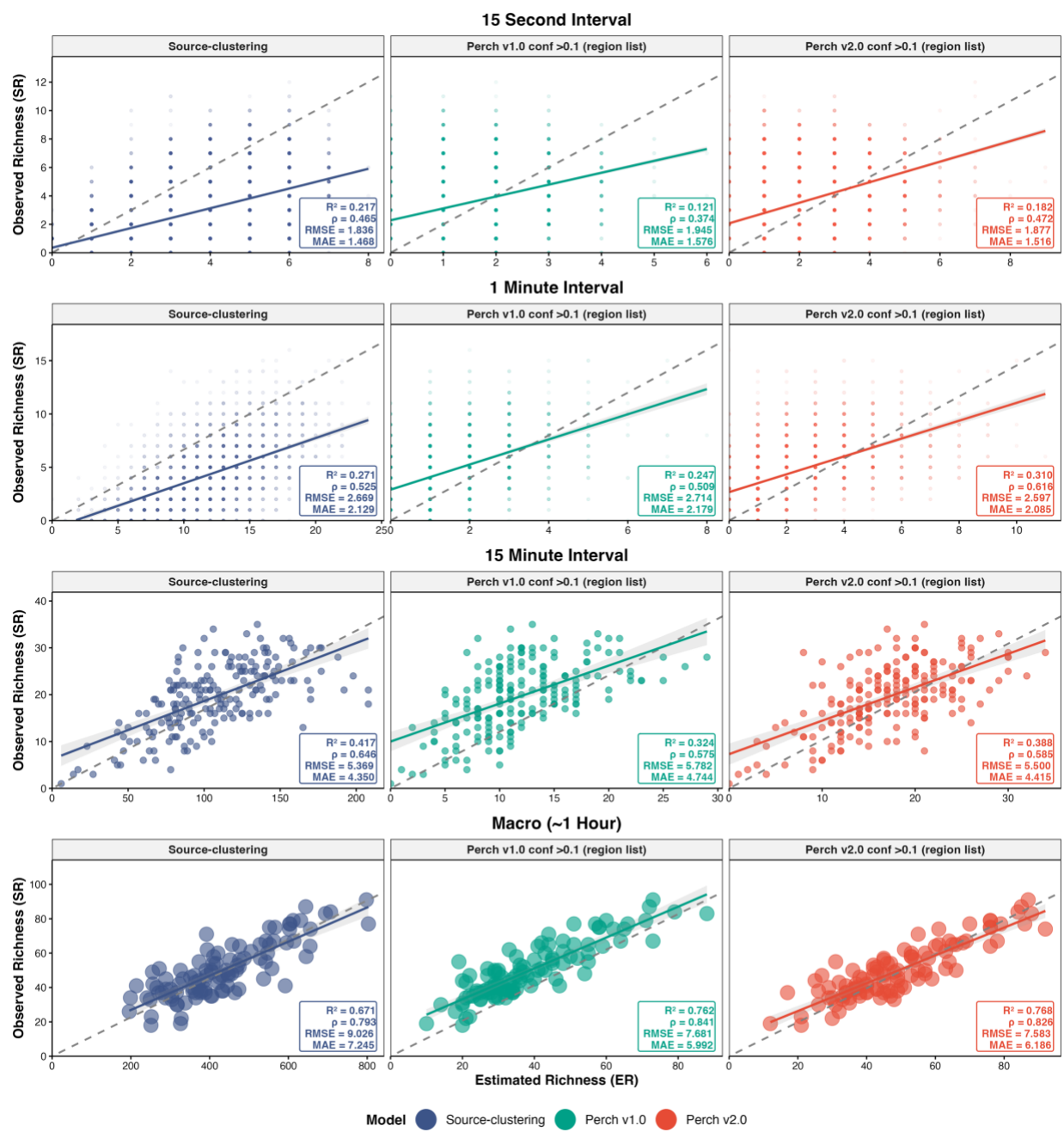
